# Lateral geniculate neurons show robust ocular dominance plasticity

**DOI:** 10.1101/097923

**Authors:** Juliane Jäpel, Mark Hübener, Tobias Bonhoeffer, Tobias Rose

## Abstract

Experience-dependent plasticity in the mature visual system is considered exclusively cortical. Using chronic two-photon Ca^2+^ imaging, we found evidence against this tenet: dLGN cells showed robust ocular dominance shifts after monocular deprivation. Most, but not all responses of dLGN cell boutons in binocular visual cortex were monocular during baseline. Following deprivation, however, deprived-eye dominated boutons became responsive to the non-deprived eye. Thus, plasticity of dLGN neurons contributes to cortical ocular dominance shifts.

---

The change in ocular dominance (OD) after monocular deprivation (MD) is one of the most prominent models of experience-dependent plasticity in the neocortex. Whereas MD is known to evoke an OD shift in the binocular part of primary visual cortex (V1)^1–4^, robust OD changes in the dorsolateral geniculate nucleus (dLGN) of the thalamus have not been reported so far^2,5^. This and similar findings led to the prevailing view that after an early developmental phase of subcortical activity-dependent changes^6^, plasticity in the visual system is exclusively cortical in the mature brain^7,8^. However, none of the recordings in dLGN to date have been performed chronically with single-cell resolution. Hence, changes in the eye-specific responsiveness of individual thalamic relay cells (TRCs) could have easily been missed. Furthermore, in contrast to the classical view of strict eye-specific segregation in dLGN^9–11^, some recent studies have reported that a substantial fraction of rodent TRCs is binocular^12,13^, which could provide a substrate for competitive TRC plasticity. This led us to re-address the questions of thalamic binocularity and experience-dependent OD plasticity, using chronic two-photon Ca^2+^ imaging of TRCs projecting to binocular V1.

We conditionally expressed the genetically encoded Ca^2+^ indicator GCaMP6m^14^ in the dLGN of Scnn1a-Tg3-Cre mice^15^ using adeno-associated virus^16–19^ and followed the visually evoked Ca^2+^ signals of individual TRC axonal boutons in layer 1 (L1) of binocular V1 for up to 4 weeks before and after MD (Fig. 1a-d). Our chronic bouton recordings provided a reliable and minimally-invasive longitudinal readout for the activity of TRCs in dLGN, specifically (see **Methods** and Supplementary Fig. 1,2).

**Figure 1.**
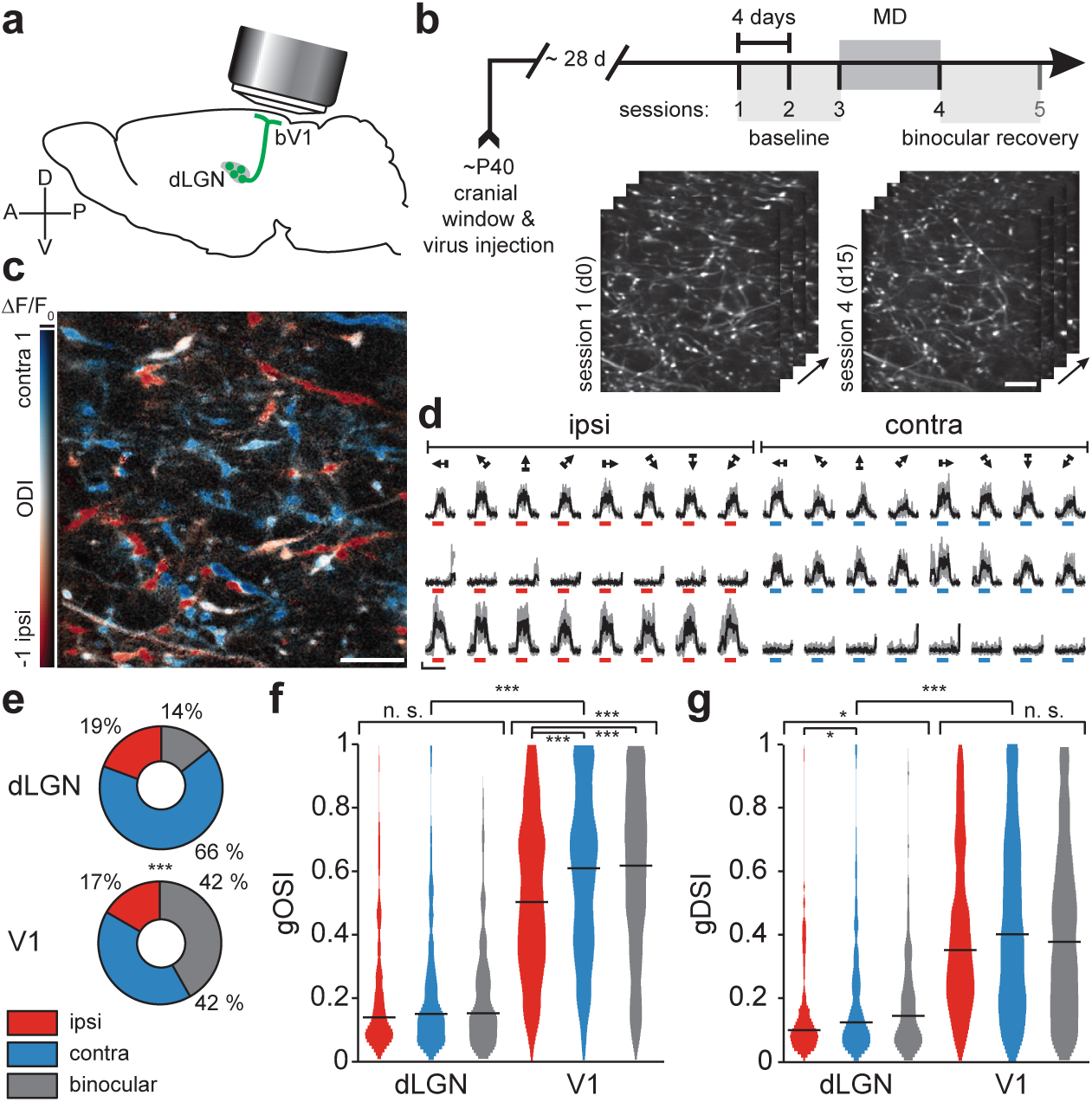
Long-term imaging and functional characterization of thalamic relay cell (TRC) boutons in L1 of mouse V1. **a)** Schematic of imaging approach. **b)** Experimental timeline and two example image volumes illustrating facilitated refinding of boutons over the extended time-course of the experiment (four slices at an image plane depth increment of 3 μm, scale bar 10 μm, frame-averaged GCaMP6m fluorescence; see **Methods** and Supplementary Fig. 1). **c)** Example of a color-coded response map of individual TRC boutons. Ocular dominance (OD) is depicted as the pixel-wise peak fluorescence change in response to ipsi- and contralateral eye preferred grating presentation (scale bar 10 μm). Red hues indicate ipsilateral dominance (ocular dominance index, ODI = (ΔF/F_contra_ – ΔF/F_ipsi_) / (ΔF/F_contra_ + ΔF/F_ipsi_); ODI < 0), and blue hues indicate contralateral dominance (ODI > 0). **d)** Ca^2+^ signals in a binocular (top) and two monocular example boutons in response to ipsi- and contralateral eye drifting grating stimulation (scale bars: ΔF/Fo = 200 %, 10 s). **e)** Fraction of responsive dLGN boutons and excitatory L2/3 cell bodies in binocular V1^4^ showing purely ipsilateral, purely contralateral or binocular significant responses (dLGN, n = 725 boutons, 9 mice; V1, n = 2.095 cells, 14 mice; P < 10^-3^, χ^2^ test). **f, g)** Violin plots of gOSI (**f**) and gDSI (**g**) of ipsi-, contralateral and binocular dLGN boutons and L2/3 cell bodies, respectively. Horizontal lines indicate medians, shaded areas the mirrored probability density estimates (dLGN, n = 725 boutons, 9 mice; V1, n = 2.095 cells, 14 mice; all gOSI dLGN vs. V1, P < 10^-193^, all gDSI dLGN vs. V1, P < 10^-120^, Mann-Whitney U-test; gOSI dLGN, P = 0.65; gDSI dLGN, P = 0.016, Kruskal-Wallis test, ipsi- or contralateral vs. binocular P = 0.11 and P = 1, Bonferroni corrected; gOSI V1, P < 10^-6^; gDSI V1, P = 0.066, Kruskal-Wallis test, ipsi- vs. contralateral or binocular gOSI P < 10^-5^ and P < 10^-6^, Bonferroni corrected; see Supplementary Fig. 3 for tuned TRC examples and further quantification). Here and in the following figures, *P <0.05, **P < 0.01, ***P < 0.001.

Recent electrophysiological studies have reported a large fraction of binocular neurons in rodent dLGN^12,13^. It is, however, difficult to rule out that this could have been the result of imperfect single unit isolation. Two-photon Ca^2+^ imaging circumvented this problem by allowing us to measure the OD of single boutons along dLGN afferents in V1 (Fig. 1c-e). We indeed observed unambiguous responses to both eyes in a subset of boutons (Fig. 1d). However, the binocularly responsive fraction (dLGN: 14%, Fig. 1e) was considerably smaller than that reported in recent electrophysiological experiments^13^. When comparing the OD of dLGN afferents with that of excitatory L2/3 cells in binocular V1, we found that, as expected, cortical neurons were significantly more binocular than dLGN afferents (cortex: 42%, Fig. 1e, P < 10^-3^), whereas the ratio of contralateral-over ipsilateral-eye population responses was comparable (Supplementary Fig. 3e,f, P = 0.36). Therefore, mouse V1 does not simply inherit binocularity from binocular geniculate cells, but, similar to carnivorans and primates^5^, predominantly integrates eye-segregated dLGN input.

We compared the stimulus selectivity of binocular TRCs to their monocular counterparts and to excitatory L2/3 cells in V1. We found, as expected^18–21^, that TRCs were far less orientation- and direction-selective than cortical neurons (Fig. 1f,g; gOSI: P < 10^-193^; gDSI: P < 10^-120^). Similar to cortex, ipsilateral TRC boutons tended to be slightly less sharply tuned than contralateral boutons (Fig. 1f,g). However, the stimulus selectivity of binocular and monocular TRC boutons did not differ in any of the parameters we tested (Fig. 1f,g; gOSI: P = 0.65; gDSI: P = 0.016, with ipsilateral and contralateral vs. binocular gDSI P = 0.11 and P = 1, respectively), arguing against the possibility that binocular dLGN cells may represent a separate visual processing channel in mice.

Does the mature dLGN show experience-dependent plasticity? To test this, we subjected adult mice to 6-8 days of contralateral eye MD (see **Methods**). Contrary to the prevailing view, we observed a clear (Fig. 2, P < 10^-8^) and largely reversible (Supplementary Fig. 4) OD shift. Whereas most TRC boutons were monocular during baseline, many acquired binocularity during MD (Fig. 2c). More than half of the responsive boutons changed their ocular dominance index (ODI) significantly (>1 standard deviation of pre-MD baseline fluctuations, Fig. 2d). In fact, the fraction of plastic boutons, the population shift magnitude, and the decrease in the fraction of responsive boutons after MD (Fig. 2e) were comparable to the effects of MD in excitatory L2/3 neurons in V1^4^ (Supplementary Fig. 5a).

**Figure 2.**
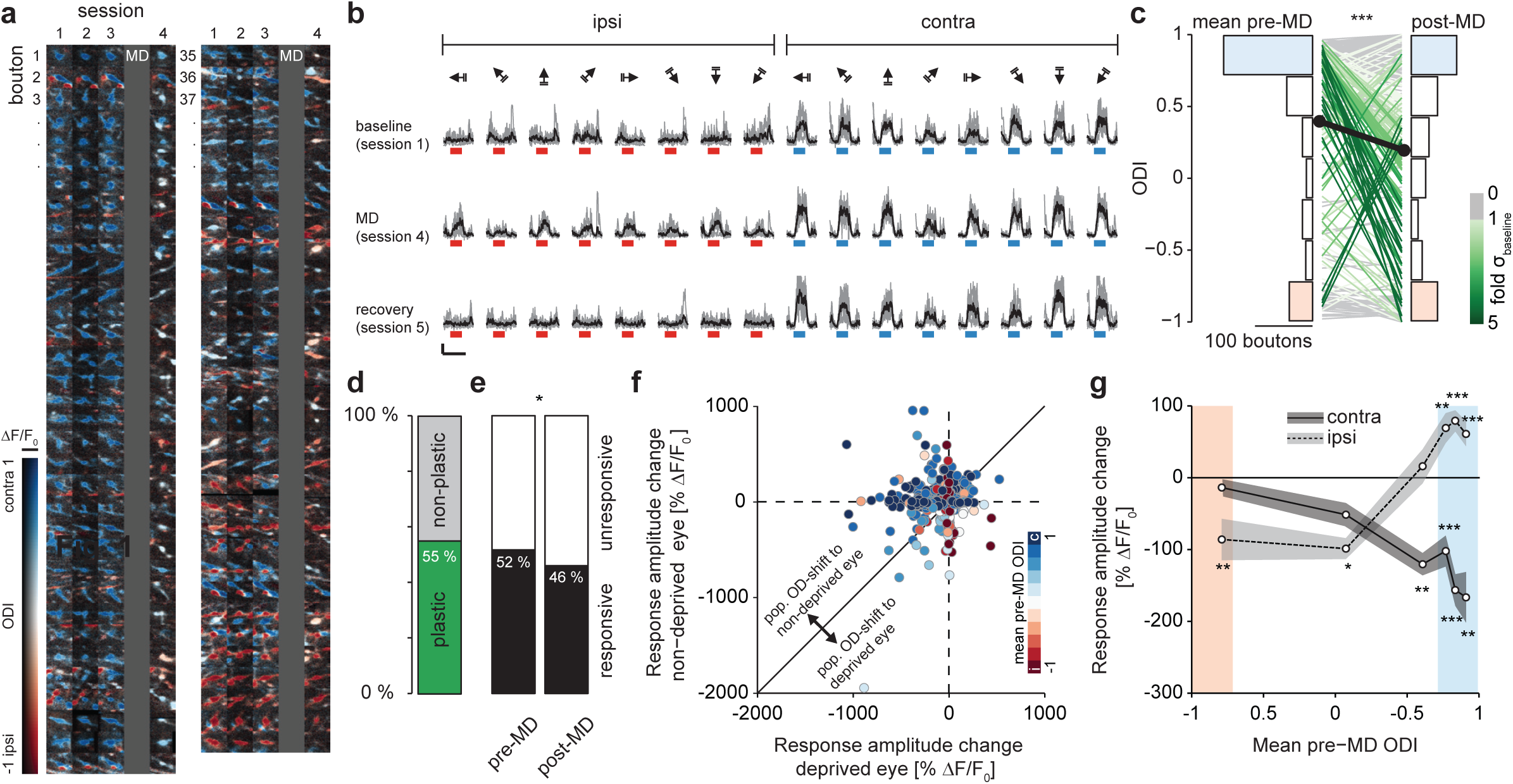
TRC boutons show robust and specific OD shifts after monocular deprivation (MD) **a)** Functional map examples of TRC boutons of a single animal (sorted by pre-MD ODI) imaged over four sessions before and after contralateral-eye MD. Note the shift from blue (contralateral) to white (binocular) and red (ipsilateral) hues after MD. **b)** Ca^2+^ signals in an example bouton in response to ipsi- and contralateral eye drifting grating stimulation over selected sessions covering OD plasticity (scale bars: ΔF/Fo = 200 %, 10 s). **c)** Single-bouton ODI distribution during baseline (mean of three pre-MD sessions, ODI = 0.40 ± 0.63, standard deviation, SD) and after 6-8 days of MD (post-MD: ODI = 0.19 ± 0.67, SD, n = 258 boutons, 9 animals, P < 10^-8^, Wilcoxon signed-rank test). Lines connect individual, continuously responsive boutons; line color indicates shift significance in units of SDs over pre-MD baseline fluctuations. Colored ODI histogram bins indicate class definitions for contralateral (blue), binocular (white) and ipsilateral (red) boutons. **d)** Fraction of boutons showing a significant (>1 SD) MD-evoked ODI change (n = 142 out of 258 boutons). **e)** Fraction of responsive and unresponsive boutons before and after MD (n = 896 boutons, P = 0. 021, χ^2^ test). **f)** Bouton-wise change in eye-specific response amplitude after MD (n = 352 boutons responsive in all pre-MD sessions, color-coded for mean pre-MD ODI). The concentration of blue data points in the upper left quadrant indicates that the population OD shift towards the non-deprived eye is largely driven by initially contralateral boutons losing contralateral-eye and gaining ipsilateral-eye responsiveness. **g)** Quantification of **(f)**, grouped into pre-MD ODI sextiles comprising similar numbers of boutons (58-59 boutons per class, paired t-tests; monocular ODI classes indicated by red and blue shading as defined in (**c**)). See Supplementary Fig. 5 for comparison with excitatory L2/3 cells in V1.

On the population level, the OD shift resulted from a net increase in ipsilateral-eye and a decrease in contralateral-eye responsiveness (Supplementary Fig. 4e). On the level of individual geniculate afferents, boutons that were exclusively responsive to the contralateral eye during baseline showed the most prominent decrease in contralateral-eye evoked activity (Fig. 2f,g). This form of deprived-eye depression is similar to our previous observation in excitatory L2/3 cells in V1^4^ (Supplementary Fig. 5b,c). However, very different from cortex, initially monocular contralateral boutons also showed the most prominent increase in ipsilateral-eye responsiveness (compare Fig. 2f,g and Supplementary Fig. 5b,c).

We found that many TRC boutons, even if they appeared monocular during baseline, received robust input from both eyes after MD (Fig. 2c). What are potential explanations for such a prominent OD change? One – in our view less likely – explanation resides in the fact that the dLGN receives substantial modulatory feedback from V1^22^ and weaker input from superior colliculus^23,24^. A decrease in cortical deprived-eye responsiveness^4^ (Supplementary Fig. 5b,c) could therefore, in principle, propagate to the dLGN and contribute to a decrease in thalamic responsiveness to the deprived eye that mirrors the cell-specific changes in L2/3 (Fig. 2f,g). We consider this less probable, however, because convergent modulatory cortical and subcortical feedback would at the same time have to elicit highly target-specific changes in non-deprived-eye responsiveness in order to lead to the specific strengthening of ipsilateral responses of initially contralaterally dominated TRCs (Fig. 2f,g). This is even more unlikely given the very different cortical changes in non-deprived-eye response strength. Here, all excitatory L2/3 cells, except initially monocular contralateral cells, increase their ipsilateral eye responsiveness after MD^4^ (Supplementary Fig. 5b,c). The, perhaps unexpected, but most parsimonious explanation is therefore, that the synapses of retinal ganglion cells (RGCs) onto TRCs are the site of change. Recent connectomics studies report a dramatic mismatch between a large structurally^25–27^ and much smaller functionally^11,28^ identified number of RGCs converging onto TRCs. We therefore speculate that unsilencing of structurally present but functionally silent ipsilateral RGC input^27,29^ may be the reason for the robust OD plasticity we observe in the dLGN.

Regardless of the precise mechanism, our results clearly show that – contrary to current models – the eye-specific responses of dLGN neurons are not rigid but undergo substantial functional OD plasticity. Given the paramount importance of this paradigm for theories of experience-dependent plasticity and the ever more widespread use of mice as model organisms, we conclude that one needs to exert caution when employing purely cortical interpretations of OD plasticity.

## Acknowledgements

We thank C. Huber, V. Staiger, F. Voss and H. Tultschin for technical assistance, J. Sigl-Glöckner for help with our transfection protocol, and P. Goltstein for programming help. This work was supported by the Max Planck Society, a Marie Curie Intra-European Fellowship to T.R., a Boehringer Ingelheim Ph.D. fellowship to J.J., and the Deutsche Forschungsgemeinschaft (grant SFB870 to T.R. and T.B.). Data and analysis codes are stored and curated at the Max Planck Computing and Data Facility Garching, Munich, Germany.

## Supplementary Methods

All experimental procedures were carried out in compliance with institutional guidelines of the Max Planck Society and the local government (Regierung von Oberbayern).

### Virus injection and chronic window implantation

Female Scnn1a-Tg3-Cre mice^1^ (postnatal day P35-45) were anesthetized by an intraperitoneal injection of a mixture of Fentanyl (0.075 mg/kg), Midazolam (7.5 mg/kg) and Medetomidine (0.75 mg/kg). Analgesia was achieved by topical application of 10% Lidocaine and subcutaneous injection of Carprofen (5 mg/kg before surgery and on the first day of post-surgical recovery). Eye dehydration was prevented by applying a thin layer of cream on both eyes. For transfection of thalamocortical boutons, the mouse was mounted in a motorized stereotactic apparatus. A circular craniotomy (4 mm diameter) was made over the somatosensory and primary visual cortex of the right hemisphere. A Hamilton Glass Syringe (gauge 36) was targeted to the dLGN (stereotactical coordinates relative to bregma: anterior 2.06 mm, lateral 2.05 mm) and a total volume of 50 to 100 nl of AAV2/1-hSYN-FLEX-GCaMP6m (1.9 × 10^13^ genome copies (GC) per ml) was injected at a depth of 2.85 mm and at a rate of 3-5 nl/s using an automated microinjector.

Imprecise virus injections could lead to two sources of contamination of our data by GCaMP-expressing axons not originating from dLGN: (i) Cortical neurons in the injection tract, and (ii) neurons of the lateral posterior nucleus (LP) of the thalamus^2–4^. We ensured that we exclusively record from dLGN afferents by several means: (i) The L4- and thalamus-specificity of the Scnn1a-Tg3-Cre driver line^1^ prevented accidental transfection of visually responsive cortical projection neurons along the injection path. (ii) We controlled for potential contamination by LP axons by post-hoc inspection of the transfection sites (see below and Supplementary Fig. 1). (iii) We measured the visual receptive field size of individual axons (Supplementary Fig. 2). Similar to previous observations^2,5^, the average receptive field size of recorded TRC boutons (area: 195 ± 88 deg^2^, median ± IQR) was slightly, but not significantly, smaller than that of excitatory L2/3 cells in V1 (Supplementary Fig. 2c) – but more than two-fold smaller than that reported for projections from LP^2,5^.

All data pertaining to experiments on excitatory layer 2/3 cells in V1 have been part of the dataset we previously obtained for Rose et al. 2016^6^. We refer to this publication for an extended description of the methods. Briefly, in this case the craniotomy was centered over the binocular zone of the right primary visual cortex and 50-200 nl of a mixture (4:1) of AAV2/1-CAG-FLEX-mRuby2-GSG-P2A-GCaMP6s-WPRE-SV40 (1.2 × 10^13^ GC/ml) and AAV2/1-CamKII0.4-Cre-SV40 (1.8 × 10^13^ GC/ml) was injected at a depth of 200–500 μm at 1-3 sites using glass pipettes and a pressure micro-injection system.

In all experiments, a cover slip (4 mm) was placed over the craniotomy and sealed with cyanoacrylate glue, and a custom-made aluminium head-plate was fixed to the skull with dental cement. Surgical anaesthesia was counteracted by subcutaneous injection of Nalaxone (1.2 mg/kg), Flumazenil (0.5 mg/kg) and Atipamezole (2.5 mg/kg). All viruses were obtained from the vector core of the University of Pennsylvania Gene Therapy Program.

### Monocular deprivation

Mice underwent monocular deprivation (MD) at the age of P70-90 as described previously^6,7^. Briefly, anaesthesia was induced by intraperitoneal injection of Fentanyl (0.05 mg/kg), Midazolam (5 mg/kg) and Medetomidine (0.5 mg/kg). Eye lids were sutured with three mattress stitches using 6-0 silk. Anaesthesia was counteracted with a mixture of Nalaxone (1.2 mg/kg), Flumazenil (0.5 mg/kg), and Atipamezol (2.5 mg/kg). After 6-8 days, mice were re-anaesthetized and sutures were removed directly before imaging. To facilitate OD plasticity in adult mice^6,8,9^, animals were housed together with at least three littermates in an enriched environment (mouse house, nesting material, running wheels) in large cages (1500 cm^2^ floor area). High contrast moving grating stimulation (8 directions) was employed for 6 h/d during the light period (14 h/d)^6,9^. Mice with corneal injuries were omitted from experiments.

### Intrinsic optical imaging

Reflected light optical imaging of intrinsic signals was used to monitor the response strength to stimulation of either eye in the binocular region of the visual cortex as a readout for ocular dominance (OD)^7^. The optical axis was adjusted to be orthogonal to the imaging window for each animal. The surface of the brain was side-illuminated with light of a 530 nm LED to visualize the blood vessel pattern and with light from two sides using a 735 nm LED (bandpass filtered at 700/40 nm) for intrinsic signal imaging. Images were collected with a CCD camera (12 bit, 260 × 348 pixel, 40Hz), which was focused 400-450 μm below the pial surface through a 4× air objective (NA 0.28). Acquisition and analysis software were custom written in Matlab. Images were clipped (1.5%) and high-pass-filtered to calculate blank-corrected image averages for each condition. These maps were thresholded (image background mean + 4 standard deviations) and the largest object was defined as the responsive region of interest (ROI). The contralateral ROI was derived from the ipsilateral ROI of the same stimulus. The mean background value of the non-responsive image region was subtracted from each pixel and all pixel values within the responsive ROI were summed to yield an integrated measure of response strength. The ocular dominance index (ODI) was determined as: *I* = 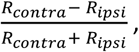 where R_contra_ and R_ipsi_ are the integrated response strengths to contralateral or ipsilateral eye stimulation, respectively.

### Chronic *in vivo* two-photon imaging

*In vivo* two-photon imaging was performed as decribed earlier^6^. Briefly, GCaMP6m was excited by a pulsed (80 MHz) femtosecond Ti:Sapphire laser (tuned to 940 nm). Signals were detected through a 16 × (NA 0.8) objective using GaAsP photomultiplier tubes (short-pass filtered at 720 nm and band-pass filtered at 525/50 nm). For imaging of thalamocortical boutons, a 57 × 57 μm field of view was scanned at 30 Hz with an 8 KHz resonant scanner at 512 × 512 pixel resolution. While imaging, the objective was rapidly moved in the z-axis by a high-load piezo z-scanner to acquire four subsequent image planes, each separated by 3 μm in depth (effective frame rate of 7.6 Hz). Volumetric multi-plane imaging was used to facilitate refinding of the same boutons across multiple imaging sessions to compensate for minor changes in brain morphology occurring during the four weeks of recording (Supplementary Fig. 1b). The average power for imaging was less than 50 mW (below the objective). For each recording session, the imaging position was realigned in x,y,z relative to the initial recording session. ScanImage 4.2^10^ and custom written hardware drivers were used for data acquisition.

Experiments were performed under light anesthesia. Mice were initially anaesthetized with an intraperitoneal injection of Fentanyl (0.035 mg/kg), Midazolam (3.5 mg/kg) and Medetomidine (0.35 mg/kg), and additional anesthetics (25% of induction level) were injected subcutaneously every 60 min to maintain anaesthesia level. Mice were placed on a heated blanket under the microscope to ensure thermal homeostasis.

### Visual stimulation

Visual stimulation was performed as described previously^6^. Briefly, MATLAB and the Psychophysics Toolbox^11,12^ were used to generate visual stimuli, which were presented on a gamma-corrected LCD. Screen position was adjusted for each mouse so that it was centered to the binocular visual field of the mouse (-15° to 35° elevation and −25° to 25° azimuth relative to midline) and the same position was kept throughout all imaging sessions.

To measure OD as well as orientation and direction tuning, 8-12 directions of drifting square wave gratings with a spatial frequency of 0.04 cycles/degree and a temporal frequency of 3 cycles/s were presented in pseudorandom order to either the left or the right eye using motorized eye shutters. Gratings were shown for 5 s, followed by 6 s of a full-field gray screen. Trials were repeated 4-6 times per eye and direction.

For receptive field mapping, we used pseudorandomized sparse noise stimulation. Stimuli consisted of black or white patches (8° × 8°) displayed on an isoluminant gray background. These patches appeared for 0.5 s with an interstimulus interval of 0.1 s at 30 different positions covering a total of 40° × 48° of the central visual field of the mouse. For each eye, this stimulus block was repeated at least ten times.

To monitor OD plasticity with intrinsic optical imaging, visual stimuli consisted of drifting square wave gratings with a spatial frequency of 0.04 cycles/degree and a temporal frequency of 2 cycles/s that were presented in a defined part of the visual field. This patch had a size of 20 × 40 degree of visual angle and was presented randomly at two different positions (next to each other in the central visual field) to either the contra- or the ipsilateral eye^7^. In each trial, a blank gray screen (50% contrast) was presented for 5 s followed by 7 s of visual stimuli (8 directions, direction changed every 0.6 s). Trials were separated by 8s and the whole stimulus sequence was repeated at least 10 times per eye and patch position.

### Histology

All dLGN injection sites were verified post-hoc (Supplementary Fig. 1). Brains were fixed with 4% w/v PFA for a week at 4°C, transferred to 30% sucrose in phosphate buffered saline (PBS) and kept at 4°C for at least three days. Brains were cut in 50 μm coronal sections using a sliding microtome. Sections were incubated overnight with 1% v/v Triton X-100 and 10% normal goat serum at 4°C. Subsequently, chicken anti-GFP (Merck Millipore, 1:1.000) was applied overnight. Sections were incubated with anti-chicken Alexa488 (Thermo Fischer Scientific, 1:200) for 3h at room temperature and then counterstained for 20 min with fluorescent Nissl stain (Molecular Probes, 1:100) and DAPI (1:10.000). Images of 512×512 pixels were acquired with an epifluorescence microscope (Zeiss Axio Imager M2) using a 10x objective. If necessary, mosaic images were acquired and realigned post-hoc.

### Image analysis

Custom-written software in MATLAB and ImageJ (http://rsbweb.nih.gov/ij/) was used for image and data analysis. The average of 160 ‘dark frames’ acquired without laser excitation was subtracted from all frames to correct for non-uniform background signal. Motion artefacts were removed by translational registration of all frames based on the average of the initial 100 signal frames of the recording. ROIs corresponding to putative boutons were automatically selected using ImageJ. A global threshold was applied to an average activity map, edge artefacts were removed, and particle-ROIs were selected based on a set of morphological filters for size and shape. The resulting ROIs were visually inspected and, if necessary, artefacts were removed. ROIs were manually matched across sessions, selecting the z-plane with optimal optical sectioning of a respective bouton for each recording (Supplementary Fig. 1b).

All pixel values within an ROI were averaged to extract the fluorescence time course (F). A sample-wise F0 trace was generated for each ROI by linear extrapolation over the low-pass filtered (0.004 Hz cutoff) local minima of the raw fluorescence signal in the absence of drifting grating stimulation (pre-stimulus periods). This F_0_ trace was used to generate the ΔF/F_0_ trace which was aligned to stimulus-onsets and was further corrected for residual offset by subtracting the individual average pre-stimulus ΔF/F_0_ signal. Boutons were defined as responsive for a specific eye if a change in peak fluorescence exceeding 8 standard deviations of the pre-stimulus signal (peak ΔF/F_0, Stimulus_ > 8 * σΔF/F_0, Baseline_) was observed in at least 50% of the trials of a single simulation condition for a specific eye. The published dataset of excitatory L2/3 cells^6^ was analyzed as described previously^6^, with the exception that now the same eye-specific responsiveness criterion as for thalamocortical boutons was used. This led to minor differences in cell selection in comparison to our previous analysis (Supplementary Fig. 5, compare to Fig. 2d in Rose et al. 2016^6^).

### Data analysis

Ocular dominance was determined by calculating the ODI from the mean ΔF/F_0_ response to the preferred eye-specific grating direction:

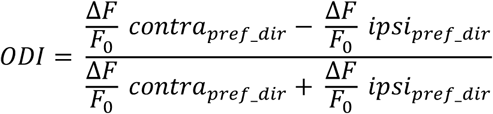

Contralateral and ipsilateral dominance are indicated by an ODI of 1 or −1, respectively. For chronic single-bouton changes in ODI, only boutons that were responsive throughout baseline and after MD (Fig. 2c), and, if applicable, during recovery (Supplementary Fig. 4f-h) were considered. For the quantification of changes in eye-specific response amplitude relative to the mean baseline ODI the 3 baseline sessions had to be responsive (Fig. 2f,g; Supplementary Fig. 5c,d). Pixel-based color-coded ODI maps were similarly generated by obtaining ODI values from the maximum stimulus evoked fluorescence change of individual pixels. Hue was set by the ODI (red: ipsilateral dominance, blue: contralateral dominance) and lightness coded the mean of the summed ipsilateral and contralateral response amplitude.

The Contra/ipsi ratio was calculated as the ratio of the mean fluorescence response to ipsilateral or contralateral eye stimulation, respectively, over all responsive boutons or layer 2/3 cell bodies of one animal (Supplementary Fig. 3e,f).

A global orientation selectivity index (gOSI) was calculated as 1 – circular variance (circ. var.)^13^:

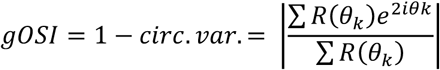

with R(θ_k_) as the mean ΔF/F_0_ response to the orientation angle θ_k_ Perfect orientation selectivity is indicated as gOSI = 1. Similarly, direction tuning was assessed by calculating the global direction selectivity index (gDSI) ^13^:

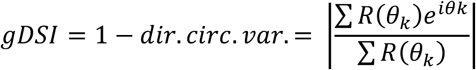

For analysis of receptive fields (RFs), traces were sorted according to stimulus position and type (black or white, representing ON and OFF subfields). Response latency was defined as the first imaging frame during a set time window in which there was a significant increase (ANOVA, p < 0.05) in fluorescence. If the ROI was determined as responsive for both the black (OFF) and white (ON) patches with different latencies, only the RF with the shorter latency was considered for further analysis. A thresholded RF subfield was derived by interpolating the raw RF at 1° resolution and only taking the region with the strongest average response. This subfield was used to compute RF area.

### Statistics

Data are reported as median ± interquartile range (IQR) or as mean ± standard error of the mean (SEM) or standard deviation (ST) as indicated in individual figures. Paired parametric (t-test) or paired and unpaired non-parametric (Wilcoxon signed-rank test, Mann-Whitney U test, Kruskal-Wallis test with Bonferroni posttest) statistics were used for comparison. For cumulative distributions, the Kolmogorov-Smirnov test was applied, while distributions were compared using χ^2^ test. Asterisks indicate significance values as follows: * p < 0.05, ** p < 0.01 and *** p < 0.001.

**Supplementary Figure 1.**
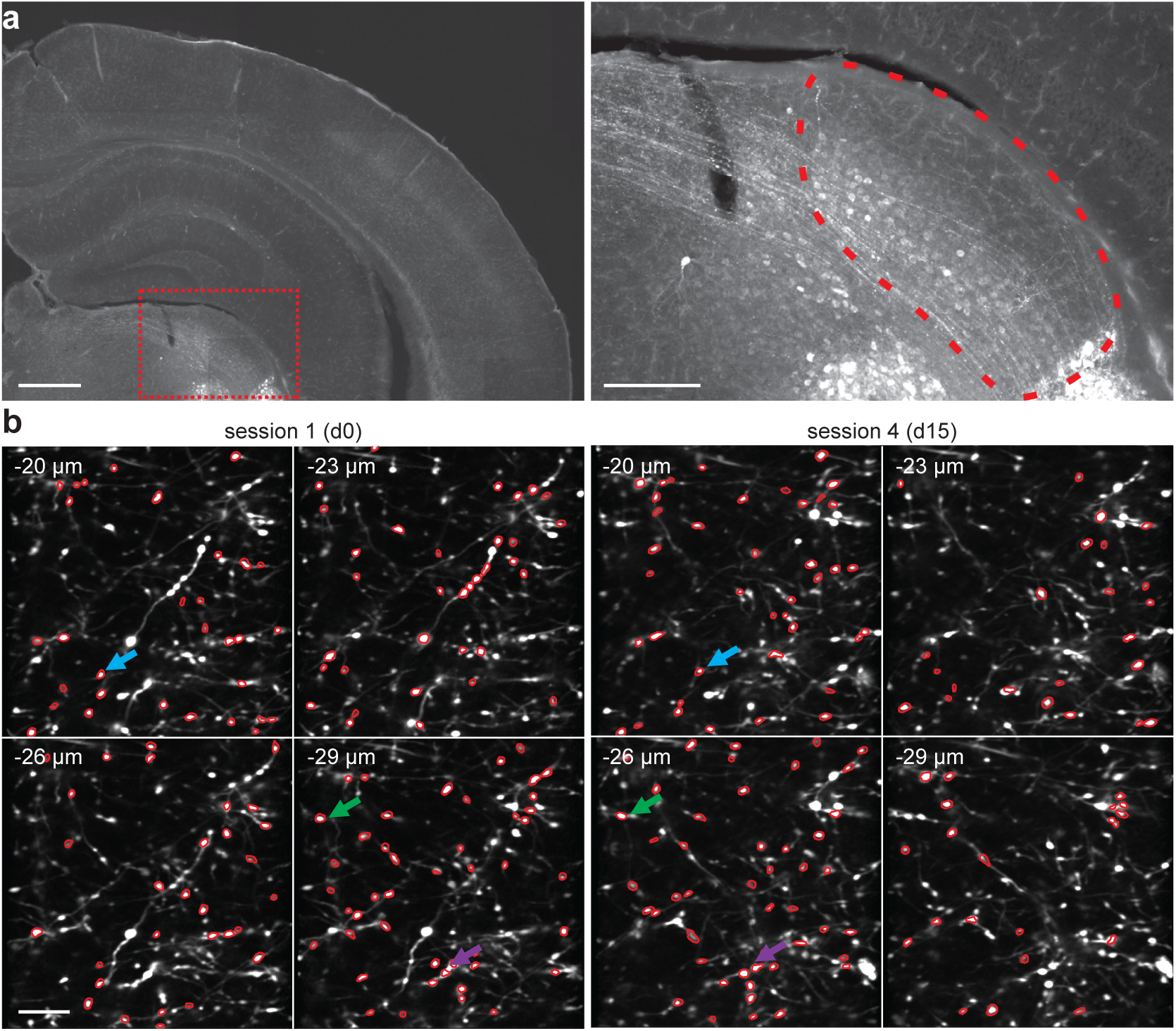
dLGN virus injections and matching of regions of interest (ROIs) across sessions **a)** AAV coding for GCaMP6m was injected into the dLGN of Scnn1A-Tg3-Cre-mice expressing Cre-recombinase in cortical layer 4 and thalamus. The representative coronal section shows GCaMP6m-labeled neurons in the dLGN (outlined in right panel, scale bar: left 500 μm, right: 200 μm). Note absence of transfected cells in the injection path and negligible dorsomedial spread into the lateral posterior nucleus (LP). **b)** Example image volumes illustrating matching of bouton ROIs across sessions. Changes in tissue morphology and positioning inaccuracies can result in individual boutons being optimally sectioned by different image planes at different time points (arrows of the same color indicate ROIs of the same group, image plane depth increment: 3 μm, first frame of volume 20 μm below pial surface; scale bar: 10 μm).

**Supplementary Figure 2.**
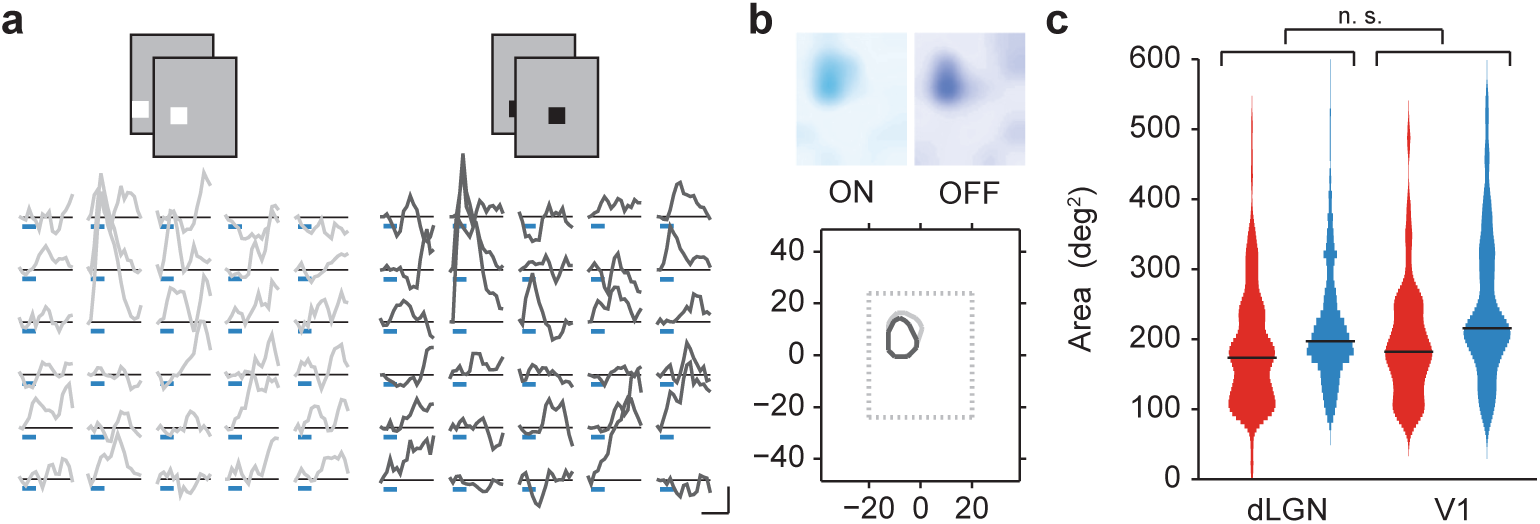
Spatial receptive field properties of thalamic relay cell (TRC) boutons in L1 of V1 **a)** Example Ca^2+^ traces (average of 10 trials) of a single bouton responding to pseudorandomized sparse noise stimulation (white, left, and black, right, patches on an isoluminant gray background) of the contralateral eye, ordered according to stimulus position (scale bars: ΔF/F_0_ = 50%, 1 s). **b)** Receptive field of the corresponding bouton in (**a**). Bottom plot shows outline of the ON and OFF sub-fields (light: ON, dark: OFF). **c**) Violin plots of spatial receptive field size of dLGN boutons and excitatory layer 2/3 cell bodies in V1 to ipsilateral (red) or contralateral (blue) eye stimulation, respectively. Horizontal lines indicate medians, shaded areas the mirrored probability density estimates (dLGN, n = 1198 receptive fields, 9 mice; V1, n = 123 receptive fields, 4 mice; combined median receptive field area dLGN: 195 ± 88 deg^2^, V1: 208 ± 99 deg^2^, P = 0.18, Wilcoxon rank-sum test).

**Supplementary Figure 3.**
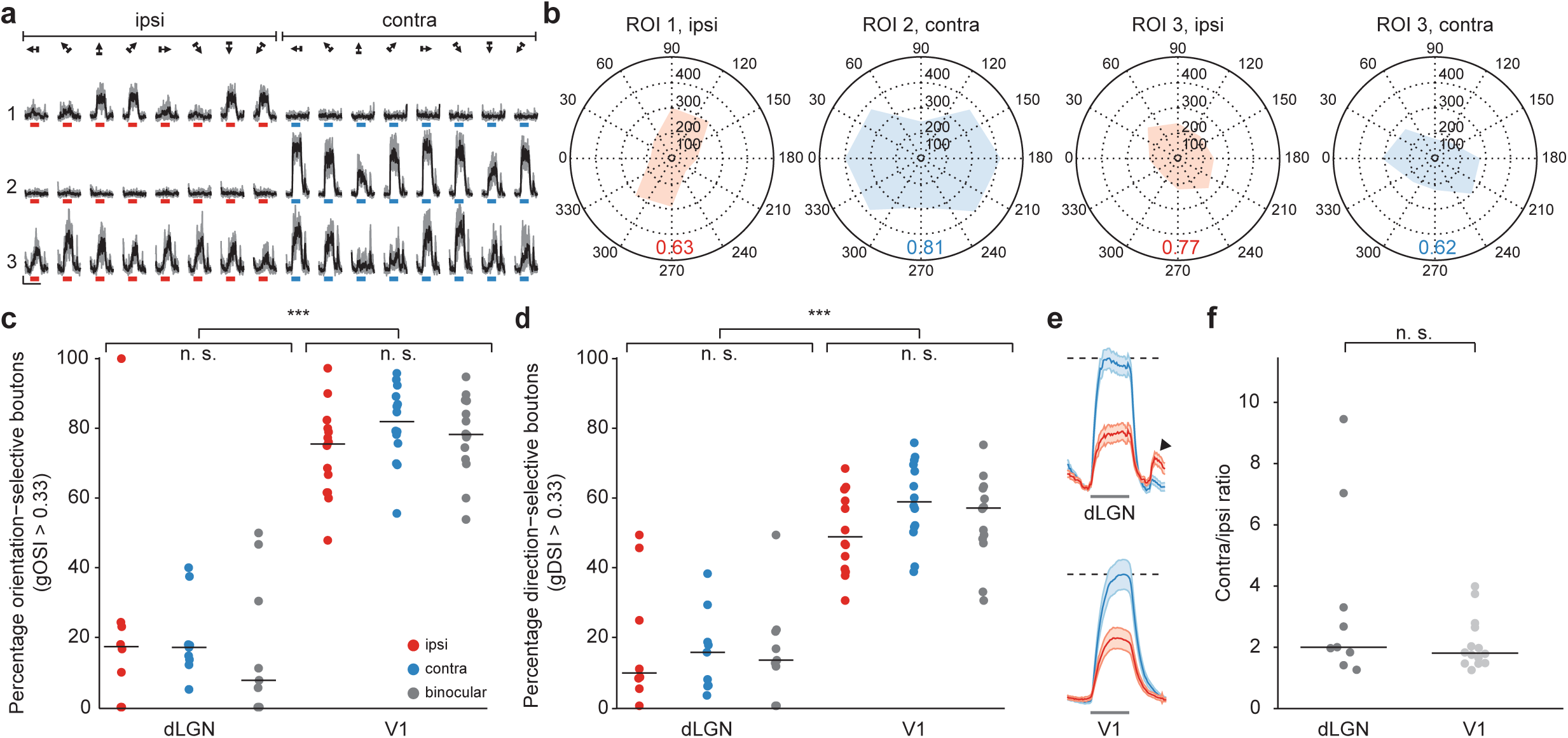
Population orientation- / direction-selectivity and eye-specific responsiveness of TRC boutons in L1 of V1. **a)** Ca^2+^ signals of three orientation-selective example boutons in response to ipsi- and contralateral eye drifting grating stimulation (scale bars: ΔF/F_0_ = 200%, 10 s). **b)** Polar plots displaying response amplitude (in ΔF/F_0_) of the boutons shown in (**a**). Numbers indicate global orientation selectivity indices (gOSI = 1 – circular variance). **c, d)** Fraction of orientation- (**c,** gOSI > 0.33) and direction-selective (**d,** gDSI > 0.33) ipsi-, contralateral and binocular dLGN boutons and V1 layer 2/3 cell bodies, respectively (dLGN, n = 9 mice; V1, n = 14 mice; all gOSI dLGN vs. V1, P < 10^-10^, all gDSI dLGN vs. V1, P < 10^-10^, Mann-Whitney U-test; gOSI dLGN, P = 0.62, gDSI dLGN, P = 0.99, gOSI V1, P = 0.20, gDSI V1, P = 0.14, Kruskal-Wallis test). **e)** Normalized contra- (blue) and ipsilalateral (red) population response of dLGN boutons (upper traces) and excitatory L2/3 cell bodies in V1 (lower traces; preferred drifting grating stimulation, normalized to peak contralateral response, dashed lines; dLGN, contra/ipsi = 2.3, n = 725 boutons; V1, contra/ipsi = 2.0, n = 2095 cells).Arrowhead indicates the pronounced dLGN on-response to pseudorandom eye-shutter switches, which is not present in cortex. **f)** Population contra/ipsi ratio in dLGN and V1 (dLGN, n = 9 mice, V1, n = 14 mice, P = 0.36. Wilcoxon rank-sum test).

**Supplementary Figure 4.**
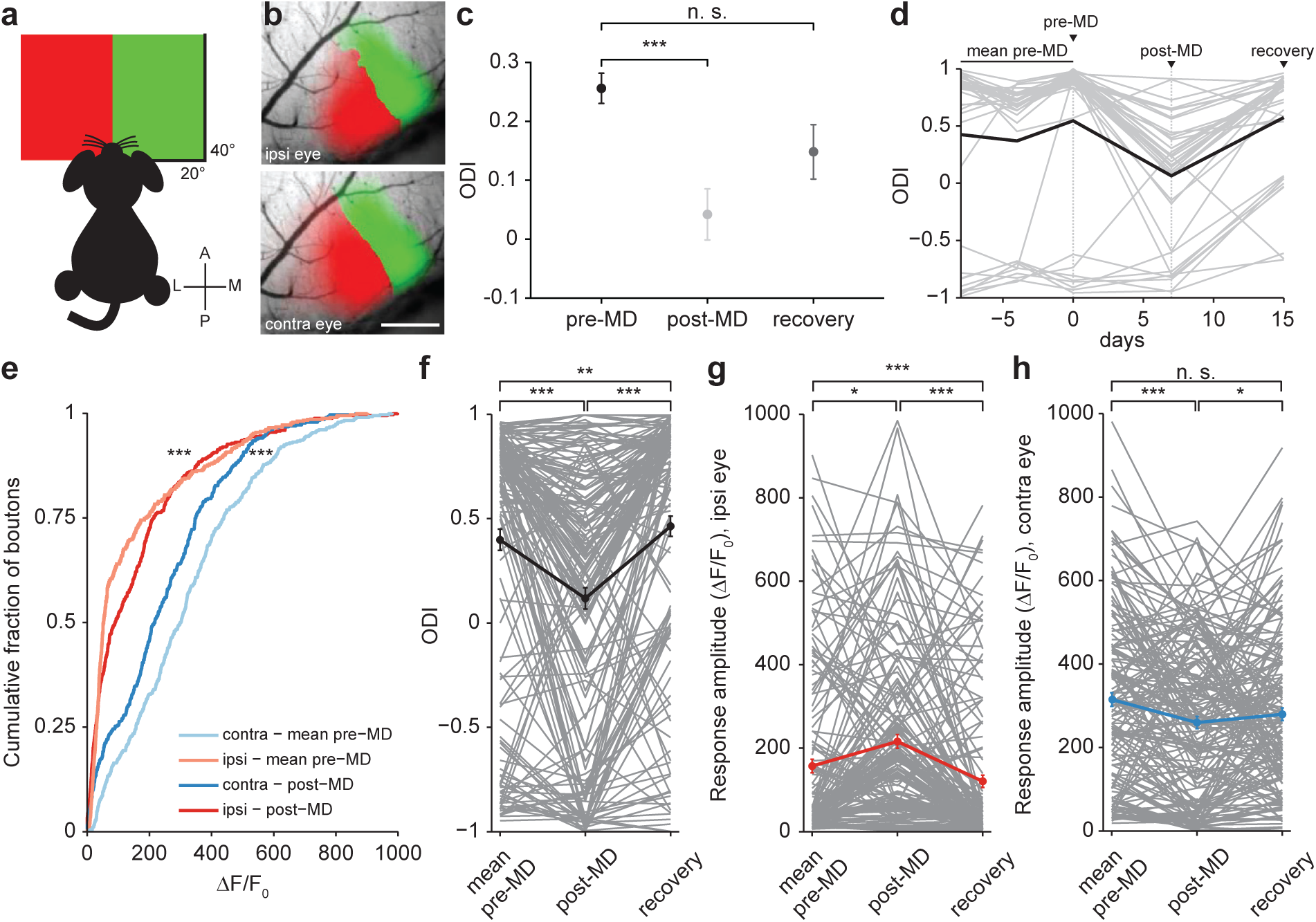
OD plasticity and recovery from OD plasticity in TRC boutons. **a)** Schematic of stimulus presentation for intrinsic optical imaging. Visual stimuli were presented at two different spatial locations on a screen in front of the mouse. **b)** Color-coded maps of cortical responses to stimulation of the ipsilateral (upper panel) or contralateral (lower panel) eye (scale bar: 0.5 mm), overlaid on an image of the cortical surface blood vessel pattern. **c)** ODI values based on intrinsic optical imaging of cortical responses for normal adult mice (mean ± SEM: 0.26 ± 0.03, n =9), after 6-8 days of MD (0.04 ± 0.04, n = 9), and after 1-2 weeks of recovery (0.15 ± 0.05, n = 6 animals, P < 10^-4^, Kruskal-Wallis test with Bonferroni's post hoc test for multiple comparisons, baseline vs. MD P = 0.0006, baseline vs. recovery P = 0.45, MD vs. recovery P = 0.41). **d)** Example time course of dLGN single-bouton ODI (gray lines) from one animal over baseline, contralateral eye MD and recovery (black line: mean ODI). **e)** Cumulative eye-specific response amplitude before and after MD (n = 456 dLGN boutons, responsive during baseline and post-MD, ipsilateral eye responses baseline vs. post-MD P < 10^-5^, contralateral eye responses baseline vs. post-MD P < 10^-6^, two-sample Kolmogorov-Smirnov tests). **f)** ODI of boutons (n = 168 boutons from 5 mice, responsive during baseline, MD and recovery) after MD and at least one week of binocular vision (P < 10^-16^, baseline vs. MD P > 10^-6^, baseline vs. recovery P = 0.002, MD vs. recovery P< 10^-10^, Friedman’s test with Bonferroni's post hoc test for multiple comparisons). **g)** Response amplitude to ipsilateral eye stimulation after MD and at least one week of binocular vision (P < 10^-13^, baseline vs. MD: P = 0.01, baseline vs. recovery P < 10^-6^, MD vs. recovery P < 10^-13^, Friedman’s test with Bonferroni post hoc for multiple comparisons). **h)** Response amplitude to contralateral eye stimulation after MD and at least one week of binocular vision (P < 10^-6^, baseline vs. MD P < 10^-6^, baseline vs. recovery P = 0.055, MD vs. recovery P = 0.035, Friedman’s test with Bonferroni post hoc for multiple comparisons).

**Supplementary Figure 5.**
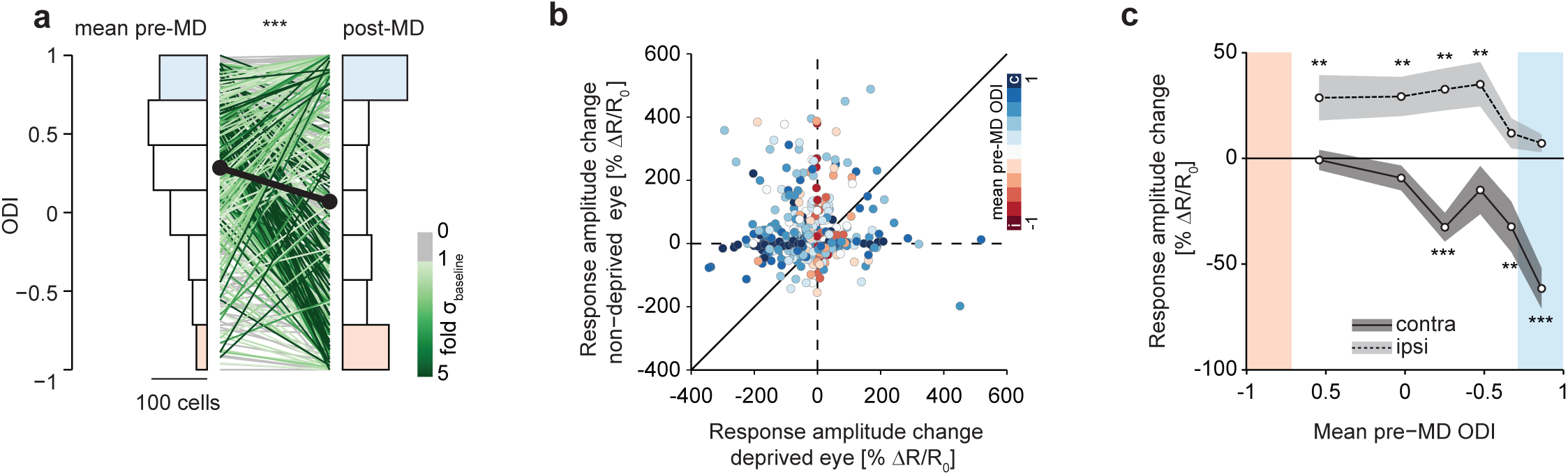
Cell-specificity of OD plasticity in excitatory layer 2/3 cells in V1. **a)** ODI distribution of the same excitatory L2/3 cells in V1 during baseline (mean of three pre-MD sessions, 0. 28 ± 0.47, SD) and after 5-8 days of MD (0.07 ± 0.67, SD, n = 430 cells in 10 mice, P < 10^-11^, Wilcoxon signed-rank test). Lines connect individual, continuously responsive cells; line color indicates shift significance in units of standard deviations over pre-MD baseline fluctuations. Colored ODI histogram bins indicate class definitions for contralateral (blue), binocular (white) and ipsilateral (red) boutons (data reanalyzed from Rose et al. 2016^6^). **b)** Change in eye-specific response amplitude after MD for individual cells (n = 565 cells responsive in all pre-MD sessions, 10 animals). Cells are color-coded by their mean pre-MD ODI. **c)** Quantification of **b)**, grouped into pre-MD ODI sextiles comprising similar numbers of cells (94-95 cells per group, paired t-tests; monocular ODI classes indicated by red and blue shading as defined in (**a**) and Fig. 2c,g).

